# Microbial Mat Functional and Compositional Sensitivity to Environmental Disturbance

**DOI:** 10.1101/063370

**Authors:** Eva C. Preisner, Erin B. Fichot, R. Sean Norman

## Abstract

The ability of ecosystems to adapt to environmental perturbations depends on the duration and intensity of change and the overall biological diversity of the system. While studies have indicated that rare microbial taxa may provide a biological reservoir that supports long-term ecosystem stability, how this dynamic population is influenced by environmental parameters remains unclear. In this study, a microbial mat ecosystem located on San Salvador Island, The Bahamas was used as a model to examine how environmental disturbance affects the activity of rare and abundant archaeal and bacterial communities and how these changes impact potential biogeochemical processes. While this ecosystem undergoes a range of seasonal variation, it experienced a large shift in salinity (230 to 65 g kg^−1^) during 2011-2012 following the landfall of Hurricane Irene on San Salvador Island. High throughput sequencing and analysis of 16S rRNA and rRNA genes from samples before and after the pulse disturbance showed significant changes in the diversity and activity of abundant and rare taxa, suggesting overall functional and compositional sensitivity to environmental change. In both archaeal and bacterial communities, while the majority of taxa showed low activity across conditions, the total number of active taxa and overall activity increased postdisturbance, with significant shifts in activity occurring among abundant and rare taxa across and within phyla. Broadly, following the post-disturbance reduction in salinity, taxa within Halobacteria decreased while those within Crenarchaeota, Thaumarchaeota, Thermoplasmata, Cyanobacteria, and Proteobacteria, increased in abundance and activity. Quantitative PCR of genes and transcripts involved in nitrogen and sulfur cycling showed concomitant shifts in biogeochemical cycling potential. Post-disturbance conditions increased the expression of genes involved in N-fixation, nitrification, denitrification, and sulfate reduction. Together, our findings show complex community adaptation to environmental change and help elucidate factors connecting disturbance, biodiversity, and ecosystem function that may enhance ecosystem models.

## Introduction

Many ecosystems are in a continuous state of change due to diel, seasonal, and intermittent extreme weather driven fluctuations in abiotic factors (e.g., nutrients, pH, light, temperature, and salinity). The ability of ecosystems to adapt to these changes depends on the duration and intensity of change and the biological diversity of the system (Sousa, 1984; Glasby and Underwood, 1996; Scheffer *et al.*, 2001; Fraterrigo and Rusak, 2008). The greatest biological diversity often exists within microbial communities involved in foundational ecosystem processes; therefore, understanding the complex links between environmental change and microbial diversity is essential for assessing ecosystem stability. Studies examining these links within lake (Shade *et al.*, 2011, 2012), marine sediment (Mohit *et al.*, 2015), soil (Allison and Martiny, 2008), and microbial mat (Boujelben *et al.*, 2012) ecosystems have shown that microbial communities are seasonally variable and often show long-term resilience to larger environmental disturbances. Further insight into the complex dynamics and ecological mechanisms maintaining ecosystems has revealed that while much of the microbial biomass is contained within a few dominant taxa, the greatest genetic diversity and potential is maintained within a vast number of low abundance or ‘rare’ taxa (Sogin *et al.*, 2006). Studies examining the significance of these rare taxa have shown that while some are highly active (Campbell *et al.*, 2011) and contribute to important biogeochemical processes (Pester *et al.*, 2010), others are less active and may be providing a genetic reservoir (*i.e.*, seedbank) to maintain ecosystem diversity over time (Jones and Lennon, 2010; Lennon and Jones, 2011). For instance, resuscitation of rare taxa has been suggested to contribute to soil ecosystem functioning (Aanderud *et al.*, 2015). However, the links between shifting environmental parameters and regulation of this dynamic population are unclear and an area of more recent research.

Complex semi-closed organo-sedimentary microbial mat ecosystems contain tightly coupled biogeochemical processes along a narrow vertical chemical gradient (Canfield and Des Marais, 1993) allowing them to serve as unique model systems to examine how environmental parameters influence the abundance and activity of rare taxa and biogeochemical cycling. While specific responses of the microbial mat rare biosphere to environmental change have yet to be explored, studies have revealed complex community dynamics with taxon-specific responses at different salinities. For instance, Cyanobacteria were shown to be tolerant to a range of salinities (Green *et al.*, 2008), while anoxygenic phototrophs grow optimally at 100-120 g kg^−1^ and sulfate reducing bacteria poorly adapted to salinities over 200 g kg^−1^ (Sorensen *et al.*, 2004). With processes within these ecosystems tightly coupled, salinity-driven shifts in community structure will alter biogeochemical cycling (Ley *et al.*, 2006). For instance, increased salinity decreases nitrogen fixation by Cyanobacteria but increases fixation by Deltaproteobacteria (Severin *et al.*, 2012). While studies have shown that microbial mat communities contain similar phyla with slight biogeographical and salinity-driven compositional differences (Ley *et al.*, 2006; López-López *et al.*, 2013; Schneider *et al.*, 2013; Allen *et al.*, 2009; Dillon *et al.*, 2013), there is little knowledge regarding the complex compositional and functional dynamics occurring within abundant and rare taxa following an environmental disturbance and how these community changes may alter biogeochemical cycling.

This study provides knowledge on microbial mat functional and compositional sensitivity following a pulse disturbance resulting in a large salinity shift. While studies have shown that salinity is one of the most important parameters influencing global patterns of Archaea and Bacteria distribution (Lozupone and Knight, 2007; Auguet *et al.*, 2010; Canfora *et al.*, 2014), these studies often compare communities isolated and adapted to a range of stable salinities. The mat ecosystem investigated here experiences routine seasonal shifts in salinity and extreme disturbances following hurricanes and tropical storms, providing an opportunity to examine the response of a semi-closed microbial community to environmental disturbance. Sequencing of archaeal and bacterial 16S rRNA and rRNA genes showed significant shifts in the day/night activity of abundant and rare taxa following the reduction in salinity from 230 to 65 g kg^−1^ that occurred between 2011-2012 following the landfall of Hurricane Irene. Quantitative PCR of genes and transcripts involved in nitrogen and sulfur cycling show concomitant shifts in gene expression indicating a possible change in biogeochemical cycling potential. Together, these data show the functional and compositional sensitivity of a microbial mat ecosystem to environmental change but also suggest that rare taxa may provide a reservoir of genetic diversity that enhances ecosystem stability following seasonal and extreme environmental disturbances.

## Materials and Methods

### Sample collection and processing

The mat ecosystem examined in this study is located on San Salvador Island, Bahamas (Supplemental Figure S1). The mat forms well-defined layers and experiences wide ranges in salinity (35 to >305 g kg^−1^), intense irradiance (> 2200 prnol m^−2^ s-^1^), and high temperatures (> 40°C) (Pinckney *et al.*, 1995; Pinckney and Paerl, 1997; Paerl *et al.*, 2003, personal observation).

Mat cores were obtained from similar locations during August 1-2, 2011 (230 g kg^−1^salinity) and 2012 (65 g kg^−1^ salinity). Samples (0.9-1.5 g) were taken during daytime (10am/5pm) and nighttime (10pm/5am) using a 7mm Harris Uni-Core^TM^ device (Ted Pella, Inc, Redding, CA, USA). Five replicate cores were pooled immediately (<1 min) into a 3 ml tube containing 2 ml RNAprotect Bacteria Reagent (Qiagen, Valencia, CA, USA). Triplicate tubes were obtained for each time point, resulting in 6 daytime and 6 nighttime replicates. Samples were homogenized with sterile glass rods and stored at 4°C until further processing. Environmental parameters were measured with an YSI 30 Salinity Meter, a YSI 55 Dissolved Oxygen Meter (YSI, Yellow Springs, OH, USA), a LI-COR LI-250A Light Meter (LI-COR, Lincoln, NE, USA), and a Mettler Toledo SevenGo Portable pH Meter (Mettler-Toledo, Columbus, OH, USA).

### Nucleic acid extraction and cDNA synthesis

Mat samples in RNAprotect Bacteria reagent were centrifuged (8,000 x g for 5 minutes) and RNAprotect discarded. The pellet was resuspended in 7.5 ml RLT Plus buffer (Qiagen) containing 1% 2-Mercaptoethanol. Samples were incubated at room temperature for 10 min followed by five freeze/thaw cycles consisting of freezing in liquid nitrogen and thawing at 55° C. Next, silicon carbide beads (DNase- and RNase-free mixture of 0.1 and 1 mm beads) were added and samples vortexed for 10 min and processed using the Allprep DNA/RNA Miniprep Kit (Qiagen) for simultaneous DNA/RNA extraction from each sample. Total RNA and DNA were quantified using a Qubit 2.0 fluorometer (Life technologies, Grand Island, NY, USA). RNA (100 ng) was reverse transcribed to cDNA using SuperScript III first-strand synthesis (Life Technologies) with random hexamers.

### 16S rRNA/rRNA gene amplification and sequencing

The abundance and activity of archaeal and bacterial communities under different salinities were assessed by sequencing and analysis of 16S rRNA and rRNA genes. For Bacteria, the V1-V3 hypervariable region was amplified using a combination (1:1 molar ratios) of the 27F forward primer (AGA GTT TGA TCC TGG CTC AG) (Edwards *et al.*, 1989) and the 27Fd forward primer (AGA GTT TGA TYM TGG CTC AG) (Nercessian *et al.*, 2005) and the U529 universal reverse primer (ACC GCG GCK GCT GRC) (Marshall *et al.*, 2012). For Archaea, the V2-V3 hypervariable region was amplified using the forward primer 109F (ACK GCT CAG TAA CAC GT) (Whitehead, 1999) and the U529 reverse primer. Twelve multiplex identifier (MID) tags were added to reverse primers for sample multiplexing (Supplemental Table S1).

Triplicate 25 μl PCR reactions contained 0.625 units of GoTaq^®^ Hot Start Polymerase (Promega Corp., Madison, WI, USA), 1.5 mM MgCl_2_, 0.2 mM nucleotide mix, 0.3 μM each primer, and 10 ng template DNA or cDNA. PCR conditions consisted of: initial denaturation: 95°C for 5:00 min; 94°C for 0:45 min, annealing at 62°C for 0:45 min using −0.5°C per cycle, and elongation at 72°C for 0:45 min, for 10 touchdown cycles; (94°C for 0:45 min, 62°C - 0.5°C per cycle for 0:45 min, 72°C for 0:45 min); 25 (for Archaea) or 15 (for Bacteria) additional cycles (94°C for 0:45 min, 57°C for 0:45 min, 72°C for 0:45 min); final elongation at 72°C for 10:00 min. Amplicons were purified using the QIAquick PCR purification kit (Qiagen) and quantified with a Qubit fluorometer.

For each sample, bacterial and archaeal amplicons with similar MIDs were combined (2:1) and uniquely labeled triplicates for each time point combined (1:1) before Illumina library preparation. Four libraries (2011 rRNA gene, 2012 rRNA gene, 2011 rRNA, 2012 rRNA) were prepared using the NEBNext^®^ DNA Library Prep Master Mix (New England Biolabs, Ipswich, MA, USA). The libraries were combined (1:1) and sequenced on an Illumina MiSeq using the MiSeq Reagent Kit v3 (Illumina, Inc., San Diego, CA, USA). FASTQ formatted paired end reads are located in the GenBank sequence read archive under SRP070186.

### Sequence processing

Sequences were analyzed using mothur [v.1.33.0; (Schloss *et al.*, 2009)] following a modified version of the MiSeq SOP (Kozich *et al.*, 2013). After paired-end reads from all libraries were assembled, sequences not matching quality criteria (maximum ambiguities = 0, ≤ 8 homopolymers, ambiguous length ≥ 300 or <650 bp) were culled using the screen.seqs command. Sequences were demultiplexed, MIDs trimmed, and separated into Archaea and Bacteria based on classification (RDPclassifier.trainset9). Non-chimeric sequences were dereplicated and aligned using Silva Archaea and Bacteria databases trimmed to the V1-V3 region. Sequences were assigned to operational taxonomic units (OTUs) at a threshold of 0.03% identity and classified. For increased stringency, OTUs represented by < 20 sequences, across groups (*i.e.*, day and night: 2011 rRNA, 2011 rRNA genes, 2012 rRNA, 2012 rRNA genes) were culled before further analysis. Sequences were rarefied (Bacteria=27,663 per MID, Archaea=24,390 per MID) and beta diversity estimated using thetaYC at a 0.03 distance threshold and visualized using non-metric multidimensional scaling (NMDS). The three dimensions of the NMDS data were plotted in SigmaPlot. Analysis of molecular variance (AMOVA) was performed to test the significance of variation among conditions.

### Activity of rare/abundant taxa

Linear discriminant analysis effect size (LEfSe, LDA>2, n=6 per condition) identified statistically significant OTUs within pre- and post-disturbance salinity conditions (Schloss *et al.*, 2009; Segata *et al.*, 2011). From initial 16S rRNA gene data, 96% of archaeal and 98% of bacterial OTUs were identified as significant among conditions and used for subsequent analysis. The relative abundance (%) of OTUs within conditions was calculated and classified as “rare” when abundance was less than 1%, and “abundant” when greater than 1%. Distribution of archaeal and bacterial OTU frequencies was explored and square root transformed to better fit a normal distribution using SPSS Statistics 22 (IBM Corp., Armonk, NY, USA). Nonparametric correlations (Kendall’s tau and Spearman’s rho) were calculated using transformed data of relative 16S rRNA and rRNA gene frequencies from 2011 and 2012 samples using SPSS and frequencies plotted with SigmaPlot. OTU activity was determined by calculating 16S rRNA:rRNA gene ratios with OTUs having a ratio above one classified as “active” and those below one classified as “slow-growing/dormant”. While this approach may not distinguish between extremely slow-growing and dormant taxa, comparison of the ratio for individual OTUs across salinities provides evidence for increased or decreased activity. Heatmaps were used to visualize OTU activities across conditions and constructed within RStudio with the heatmap2 command in gplot (Warnes *et al.*, 2012).

### Quantitative PCR

Abundance of select genes and transcripts involved in nitrogen and sulfur cycling as well as overall Archaea and Bacteria abundance was measured by quantitative PCR (qPCR). For each target, triplicate qPCR reactions were performed for each biological triplicate in an ABI 7900HT Fast Real-Time PCR system (Applied Biosystems, Carlsbad, CA, USA) with a final volume of 25 μl, including 12.5 μl PerfeCta SYBR Green SuperMix, Rox (Quanta Biosciences, Gaithersburg, MD, USA), 300 nM respective primers (Supplemental Table S2), and 10 ng DNA/cDNA. Template DNA/cDNA consisted of pooled (equal molar) day and night nucleic acids to examine overall day/night biogeochemical cycling and domain distribution during sampling years.

Data were analyzed with SPSS using an independent sample t-test (two-tailed, p<0.05) to detect differences in RNA:DNA ratios between 2011 and 2012, as well as multiple comparison analysis (ANOVA) of 16S rRNA genes to detect differences in Archaea and Bacteria abundances.

## Results and Discussion

### Post-disturbance shifts in community diversity

Following the reduction in salinity, qPCR of 16S rRNA genes from replicate day/night samples indicated significant (p<0.05) changes in overall domain prevalence with Archaea decreasing from 57% to 33% and Bacteria increasing from 43% to 67%. Further differences in community composition (β-diversity) between salinity conditions showed that, within respective years, Archaea and Bacteria day and night samples were uniformly clustered, suggesting minimal diel influence on community composition (Supplemental Figure S2). However, spatial separation was observed between years as 2011 and 2012 samples formed distinct clusters, revealing an overall shift in the archaeal and bacterial community with decreased salinity. Analysis of molecular variance (AMOVA) further supported the significant difference in community composition between years (p<0.0001). These results are similar to other studies that have found salinity to be a main determinant factor in controlling microbial diversity within soil, sediment, solar saltern, and lake ecosystems (Benlloch *et al.*, 2002; Rietz and Haynes, 2003; Tripathi *et al.*, 2005; Lozupone and Knight, 2007; Canfora *et al.*, 2014; Webster *et al.*, 2015; Aanderud *et al.*, 2016). While the distribution of active taxa (16S rRNA) was even across the archaeal community during both years, the pre-disturbance high saline conditions of 2011 selected for a distinct active bacterial population as compared to more even distribution of activity observed after the post-disturbance reduction in salinity (Supplementary Figure S2). These data suggest that the archaeal community contains habitat generalists with a wide range of osmotic tolerance while the bacterial community consists of more habitat specialists that are selected under different environmental conditions. These results support a previous study that observed selection of bacterial specialists within isolated ecosystems that have adapted to salinity extremes (Logares *et al.*, 2013).

### Post-disturbance shifts in archaeal OTU abundance

To better understand how the archaeal community shifted following the reduction in salinity, the relative abundance of archaeal 16S rRNA genes was compared across years (Figure 1A). Due to similarities in day/night abundances, average abundances are discussed for comparison between years. During both pre- and post-disturbance years, 3.5% of archaeal OTUs had relative sequence abundances greater than 1% and classified as ‘abundant’ while the majority were less than 1% and classified as ‘conditionally rare’. Although rare OTUs comprised a majority of the community richness, in total they represented only 33% and 39% of overall archaeal 16S rRNA gene abundance in 2011 and 2012, respectively. Following the reduction in salinity, OTU abundance shifted with 4 OTUs remaining abundant across salinities, while 3 shifted from abundant to rare and 11 from rare to abundant. Additionally, 168 OTUs were detected under only one salinity condition, suggesting greater environmental specialization by taxa than that suggested by the lower-resolution NMDS plots. Other studies have found similar rare/abundant distribution among microbial communities examined in a range of ecosystems, where they are suggested to provide a long-term genetic reservoir (Jones and Lennon, 2010; Lennon and Jones, 2011; Aanderud *et al.*, 2015).

**Figure 1.**
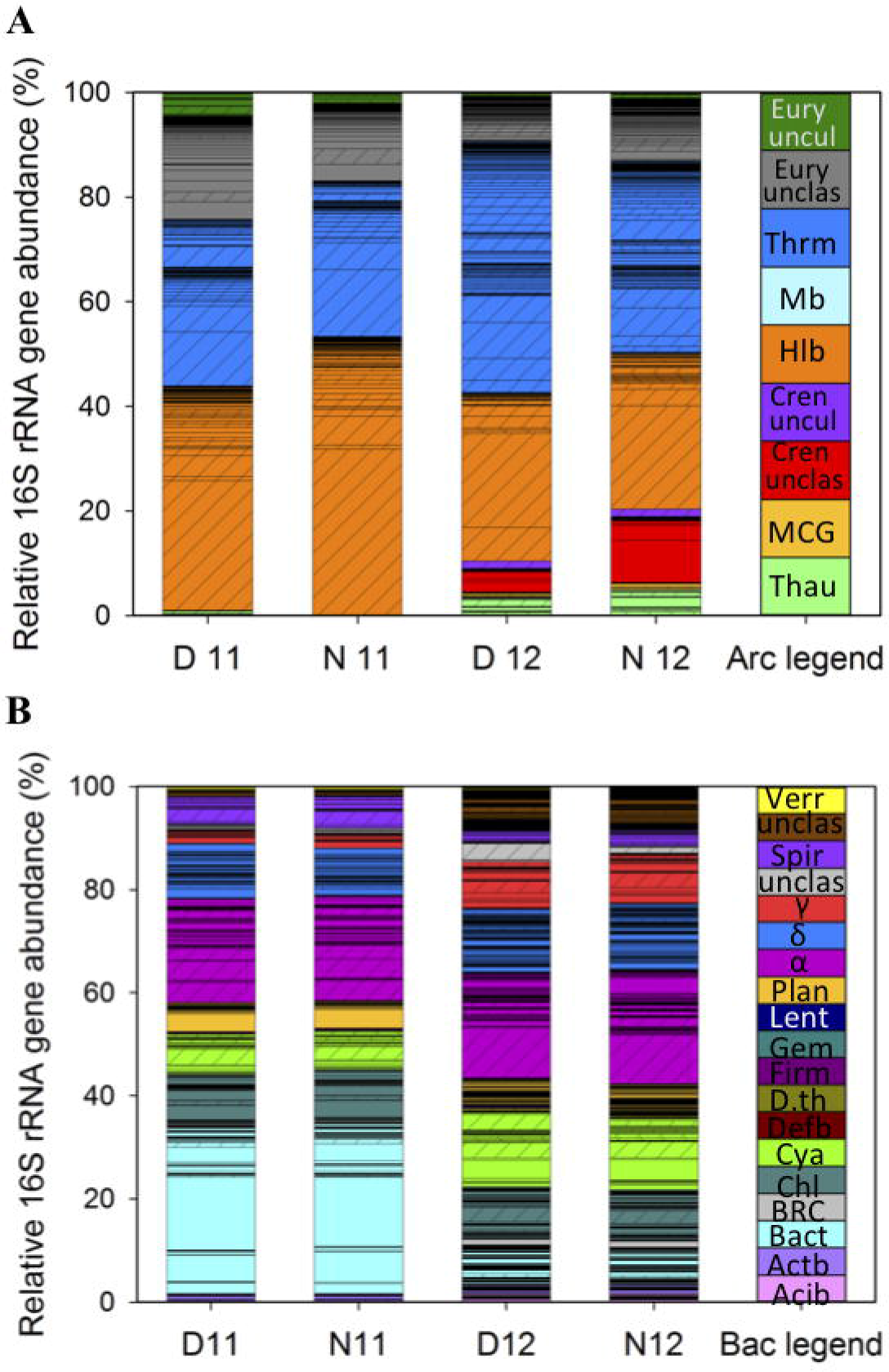
Archaea (A) and Bacteria (B) OTUs sorted by phylogenetic classification during day and night under high and lower salinity. Bars represent the average (n=6) relative 16S rRNA gene abundance of each OTU under the following conditions: D11, day 2011; N11, night 2011; D12, day 2012; N12, night 2012). Cross hatched bars represent OTUs present under both salinities. Abbreviations: **Archaea legend** Thau, Thaumarchaeota; MCG, Miscellaneous Crenarchaeotic Group; Cren unclas, Crenarchaeota unclassified; Halo, Halobacteria; Mb, Methanobacteria; Thrm, Thermoplasmata; Eury unclas, *Euryarchaeota* unclassified; Eury uncul, Euryarchaeota uncultured. **Bacteria legend** Acib, Acidobacteria; Actb, Actinobacteria; Bact, Bacteroidetes; BRC1, Bacteria Candidate division BRC1; Chl, Chloroflexi; Cya, Cyanobacteria; Defb, Deferribacteria; D.th, Deinococcus Thermos, Firm, Firmicutes; Gem, Gemmatimonadetes, Lent, Lentisphaerae; Plan, Planctomycetales; Proteo, Proteobacteria; α, Alphaproteobacteria, δ, Deltaproteobacteria; γ, Gammaproteobacteria; unclas, Proteobacteria unclassified, Spiro., Spirochaetales; unclas, Bacteria unclassified; Verr, Verrucomicrobia.

Classification of OTUs showed that under the 2011 high salt conditions, the single most abundant class was Halobacteria, representing approximately 48% of total archaeal 16S rRNA gene abundance (Figure 1A, D/N11). While most abundant, the evenness of the Halobacteria community was skewed toward a single OTU belonging to the family, Halobacteriaceae. Taxa within this physiologically diverse class are defined by their halophilic nature and have been widely observed in a number of hypersaline systems where they couple phototrophy with osmoregulation (Ochsenreiter *et al.*, 2002; Sorensen *et al.*, 2005; Maturrano *et al.*, 2006; Oren, 2008; Youssef *et al.*, 2012; Sorokin *et al.*, 2014; Williams *et al.*, 2014; Najjari *et al.*, 2015). While the single OTU discussed above remained the most abundant following the post-disturbance reduction in salinity, the total number of Halobacteria OTUs decreased from 58 to 28 resulting in a 52% reduction in richness (Figure 1A; D/N12). In addition, a 37% reduction in Halobacteria abundance was observed with taxa decreasing from 48% to 31% of the total post-disturbance archaeal community. The next most abundant classes identified under high salinity were unclassified Euryarchaeota and Thermoplasmata, which showed the highest OTU richness across years but only represented approximately 17% and 31% of overall archaeal 16S rRNA gene abundance, respectively (Figure 1A; D/N11). Following salinity reduction, previously abundant Thermoplasmata OTUs decreased while previously rare or not detected OTUs expanded, with an overall increase from 77 to 126 detected OTUs, resulting in a 63% increase in richness and a 37% increase in abundance (Figure 1A; D/N12). Increased richness and abundance suggest a wider range of Thermoplasmata taxa adapted to lower salinity. Thermoplasmata OTUs were further classified into the Thermoplasmatales order, which is comprised of thermophilic and acidophilic taxa often observed in anoxic marine environments (Ferrer *et al.*, 2011). Other studies have observed Thermoplasmata taxa within microbial mats but their role within these ecosystems is still largely unknown (Schneider *et al.*, 2013; Robertson *et al.*, 2009). However, with the identification of *mcrA* genes within some taxa identified through 16S rRNA gene studies as Thermoplasmata or uncultured/unclassified Euryarchaeota (Paul *et al.*, 2012), it is likely that many of these taxa have a methanogenic role within high salinity conditions such as those found within hypersaline microbial mats. Within the archaeal community, the greatest post-disturbance shift in abundance was observed for Crenarchaeota, with unclassified and uncultured Crenarchaeota increasing from 0.3% to 11% of the community 16S rRNA gene abundance (Figure 1A). Crenarchaeota are widely distributed in marine and lake sediments as well as microbial mats and while important in global sedimentary processes (Robertson *et al.*, 2009; Kubo *et al.*, 2012), their growth appears suppressed under extreme hypersaline conditions such as those observed during 2011. While few archaeal OTUs across salinities classified within the Thaumarchaeota phylum, an increase in richness and abundance was observed with the reduction in salinity. Taxa within this phylum are involved in ammonia-oxidation (Brochier-Armanet *et al.*, 2008; Pester *et al.*, 2011) and play important roles in nitrogen cycling within marine environments (Spang *et al.*, 2010) and microbial mats (Schneider *et al.*, 2013; Reigstad *et al.*, 2008).

Overall, the pre-disturbance conditions of 2011 favored Halobacteria, which are known to thrive in environments with salinities over 100 g kg^−1^ (Oren *et al.*, 2009), while post-disturbance conditions were selective for Crenarchaeota, Thermoplasmata, and Thaumarchaeota growth. Furthermore, similar to a study demonstrating dynamic shifts in Halobacteria taxa within aquatic systems with fluctuating salinity (Najjari *et al.*, 2015), our data show that, within classes, while many OTUs were observed across salinities, their relative abundance changed resulting in many rare/abundant shifts and suggesting class-level environmental adaptation and possible functional redundancy.

### Post-disturbance shifts in bacterial OTU abundance

To examine post-disturbance shifts in the bacterial community, the relative abundance of bacterial 16S rRNA genes was also compared across years (Figure 1B). As above, due to the similarities between abundances, day/night averages are discussed for comparison between years. Similar to the archaeal community, during both years, 1.5% of bacterial OTUs were classified as ‘abundant’ while the remaining were classified as ‘conditionally rare’. While rare OTUs comprised a majority of the community richness, in total they represented 46% and 65% of relative bacterial 16S rRNA gene abundance in 2011 and 2012, respectively. Following the reduction in salinity, OTU abundance shifted with 2 OTUs remaining abundant across both salinities, while 3 shifted from abundant to rare and 6 from rare to abundant. In addition, 403 OTUs were only detected in one salinity condition, suggesting environmental specialization by many bacterial taxa.

Classification of bacterial OTUs showed that Bacteroidetes was the most abundant pre-disturbance phylum with three OTUs classified as Sphingobacteria having the greatest abundance (Figure 1B; D/N11). Under post-disturbance conditions, while Bacteroidetes richness remained the same, a 78% reduction in relative abundance was observed with abundance decreasing from 33% to 7% within the bacterial community (Figure 1B, D/N12). This decrease suggests that the most abundant OTUs may represent halophilic taxa within Sphingobacteria, similar to a previous study showing the presence of Sphingobacteria taxa within hypersaline conditions (Schneider *et al.*, 2013). For the following phyla, Actinobacteria, Chloroflexi, Planctomycetes, and Verrucomicrobia, equal OTU richness and abundance were observed across years, however, some OTUs were only present under one salinity condition. For instance, within the phylum Planctomycetes, one OTU classified as Phycisphaerae was abundant under high salinity and decreased significantly following the shift in salinity. Extreme salt tolerance of Phycisphaerae has previously been observed in cultures isolated from hypersaline microbial mats (Spring *et al.*, 2015). Interestingly, while all Planctomycetes OTUs belonged to the rare population in 2012, over 50% were detected only at lower salinity, suggesting a shift toward more favorable growth conditions resulting in increased phylum richness.

Cyanobacteria comprised 7% of the pre-disturbance bacterial community but increased to 14% relative abundance concomitant with salinity reduction (Figure 1B). Within the Cyanobacteria, while most OTUs were classified as conditionally rare, one was abundant under both conditions while two changed from not detected or rare to abundant with salinity reduction. Cyanobacteria are often considered keystone species in many microbial mats and, due to the synthesis of numerous osmotic solutes (Oren, 2008), show a range of salt tolerance, resulting in salinity driven specialization and distribution (Braithwaite and Whitton, 1987; Dubinin *et al.*, 1992; Franks and Stolz, 2009). While a range of salt tolerance was also suggested in the current study, abundance appeared constricted at high salinity with more adaptation toward the post-disturbance lower salinity conditions of 2012.

The greatest bacterial community richness across both salinities was contained within the Proteobacteria phylum. Within this phylum, Alphaproteobacteria was the most abundant class across years, followed by Delta-, Gamma-, and unclassified-Proteobacteria. This distribution of Proteobacteria and the lack of Betaproteobacteria within the microbial mat ecosystem is consistent with other studies examining the influence of salinity on bacterial composition (del Giorgio and Bouvier, 2002; Herlemann *et al.*, 2011; Morrissey and Franklin, 2015). Furthermore, among the Alphaproteobacteria, 4 OTUs belonging to the orders Rhizobiales, Rhodobacterales, and unclassified Alphaproteobacteria were the most abundant under high salinity. Following the reduction in salinity, Rhodobacterales OTUs decreased by 42%, while a single abundant Rhizobiales OTU further increased by 155%, shifting the order to become the most abundant Alphaproteobacteria at lower salinities. While Deltaproteobacteria had the greatest richness among the Proteobacteria, with OTUs spanning 9 orders, the Desulfobacterales and unclassified Deltaproteobacteria contained the most OTUs and had the highest abundance within the class. While no change in overall abundance was observed across salinities, OTU distribution among Deltaproteobacteria changed significantly, with half of the OTUs only detected under one condition, suggesting salinity-driven succession among this population (Figure 1B, non-cross hatched OTUs). The largest shift in proteobacterial abundance across salinities was observed within the Gammaproteobacteria, which showed an increase from 2.7% to 9.1% following the decrease in salinity. This shift in abundance was correlated with an increase in three OTUs classified as Chromatiales and Sedimenticola. Chromatiales have been identified in a variety of environments such as mangroves (Gomes *et al.*, 2010), aquatic habitats (Imhoff, 2006), hypersaline lakes (Mesbah *et al.*, 2007), Salinas (Caumette *et al.*, 1988), and microbial mats (Caumette *et al.*, 1991) suggesting they may be adapted to a variety of salinities but possibly suppressed at extreme high salinities such as those observed in this study during 2011. Sedimenticola are potential denitrifiers using sulfide as electron donor (Russ *et al.*, 2014) and have been previously described in ocean sediments, indicating possible tolerance towards lower salinities.

Overall, analysis of bacterial 16S rRNA gene abundance across years showed dynamic shifts in taxa, with OTUs within the phyla Cyanobacteria, Proteobacteria, and unclassified bacteria enriched under lower salinity conditions. Given that many of these phyla are known to play a role in biogeochemical cycling, fluctuations in salinity and changes in community composition may ultimately impact overall ecosystem function. However, recent studies have shown that taxa abundance and activity are not always correlated; therefore, 16S rRNA gene abundance can not be used to solely infer community activity (Campbell *et al.*, 2011; Pester *et al.*, 2010).

### Post-disturbance shifts in archaeal OTU activity

Comparison of 16S rRNA:rRNA gene ratios was used to test the association between OTU activity and abundance during day and night across conditions (Figure 2). A positive correlation between the daytime activity and abundance of archaeal OTUs was observed during both years, suggesting an overall trend within which the activity of an OTU increased with abundance (Figure 2A; Kendall’s nonparametric τ_day2011_=0.59, p<0.0001, n=275; τ_day2012_=0.59, p<0.0001, n=330). Similarly, while a positive correlation was also observed during nighttime, a reduction in the Tau coefficient during 2012 suggested a weaker association and more variable growth under the lower salinity nighttime samples (Figure 2B; Kendall’s nonparametric τ_night2011_=0.55, p<0.0001, n=249; τ_night2012_=0.38, p<0.0001, n=367). Further analysis of archaeal OTUs under 2011 pre-disturbance daytime conditions showed that 27% of abundant OTUs (Figure 2A, red triangles right of vertical line) had 16S rRNA:rRNA gene ratios above one and classified as ‘active’ (above the 1:1 line) while the remaining abundant OTUs had ratios less than one and classified as 'slow-growing/dormant’ (below the 1:1 line). The rare population showed similar trends with 16% of archaeal conditionally rare OTUs classified as active while the majority remained slow-growing/dormant (Figure 2A, red triangles left of vertical line). While the overall trend of most archaeal OTUs being classified as slow-growing/dormant was retained after the salinity shift, the total number of active OTUs increased as compared to 2011 with a 35% and 15% increase in active abundant and rare archaeal OTUs, respectively. Similar trends were observed during nighttime (Figure 2B) with the majority of rare OTUs being slow-growing/dormant but active OTUs increasing by 10% in 2012. The increase in active OTUs during 2012 suggests that the reduced salinity provided environmental conditions more conducive to overall archaeal activity.

**Figure 2.**
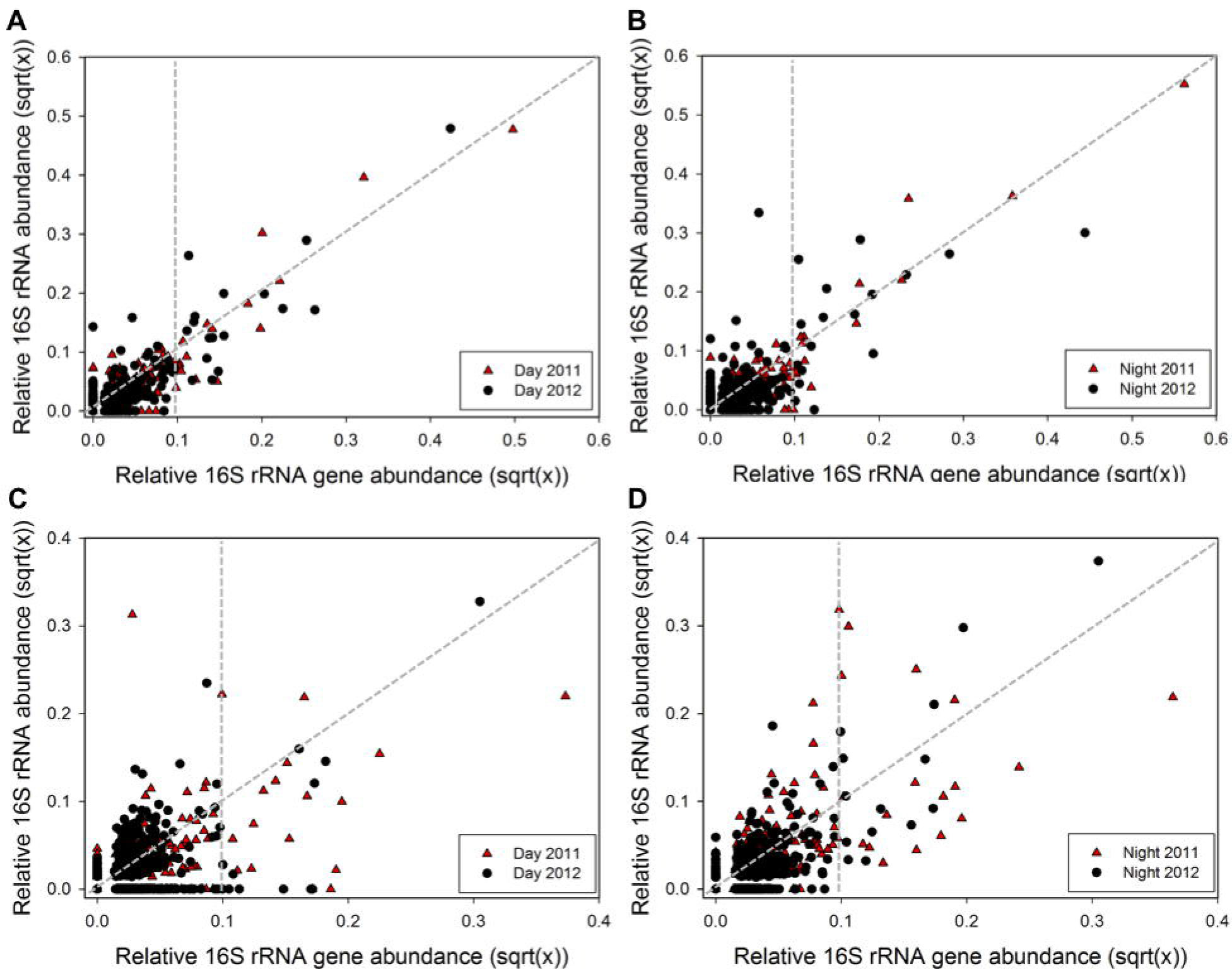
Relationship between 16S rRNA and 16S rRNA gene abundance for each OTU for subsampled communities comparing 2011 (black) and 2012 (red). Archaeal **(A)** day (n_day2011_=277, n_day2012_=3 3 0) and **(B)** night (n_night2011_=249, n_night2012_=367); and bacterial **(C)** day (n_day20111_=372, n_day2012_=697) and **(D)** night (n_night2011_=347, n_night2012_=621). Data points in graph are square root transformed paired relative 16S rRNA and 16S rRNA gene abundances for each OTU. Vertical grey dashed line differentiates between rare and abundant OTUs (rare <0.1%, abundant >0.1% 16S rRNA gene abundance). Data points above and below the 1:1 line represent active and slow-growing/dormant OTUs, respectively.

To further examine how community activity was affected by pulse disturbance, archaeal OTUs were classified and 16S rRNA:rRNA gene ratios compared during day/night across salinity conditions (Figure 3). While under both salinity conditions the highest percentage of active OTUs was contained in the rare population, the contribution of the abundant taxa may be higher due to greater overall abundance. During the pre-disturbance conditions of 2011, the greatest contribution to overall archaeal community activity was observed in OTUs within unclassified Euryarchaeota, followed by Thermoplasmata, Halobacteria, uncultured Euryarchaeota, and Methanobacteria (Figure 3, D/N11). Diel comparison at high salinity showed differences in OTU activity within most classes/phyla with a 29% and 196% increase in Halobacteria and Thaumarchaeota activity at night and increased unclassified Crenarchaeota activity during daytime (Figure 3, D vs N 11). Following salinity reduction, a shift was observed in the active archaeal population, concomitant with changes in abundance (Figure 1A), which resulted in more even distribution of activities across classes (Figure 3, D12 and N12). For instance, decreased activity was observed within unclassified and uncultured Euryarchaeota, and Halobacteria, while unclassified Crenarchaeota, Thermoplasmata, Thaumarchaeota, and miscellaneous crenarchaeotic group increased in activity, resulting in a 33% increase in overall archaeal total activity as determined by the sum of day/night 16S rRNA:rRNA gene ratios for both salinity conditions. While halobacterial OTU richness and activity decreased following the reduction in salinity, many rare OTUs increased in activity, further suggesting environmental adaptation to a range of salinities and consistent with previous study identifying dynamic shifts in Halobacteria (Najjari *et al.*, 2015). The >95% increase in abundance and activity of OTUs within the unclassified Crenarchaeota suggests that high salinity provided an environmental constraint to the growth and activity of this taxon which was relaxed following the reduction in salinity. While Crenarchaeota have been mostly observed in freshwater ecosystems (Auguet *et al.*, 2010) our results are consistent with another study that observed high altitude lake sediments of intermediate salinity dominated by Crenarchaeota while high salinity sediments were dominated by Halobacteria (Liu *et al.*, 2016). Opposite from the high salinity condition, diel trends at lower salinity showed that Halobacteria activity decreased by half while unclassified Crenarchaeota and Thermoplasmata activities increased at night. Overall, the influence of salinity on Archaea activity is complex with environmentally-driven adaptation and dynamic shifts not only across but also within classes.

**Figure 3.**
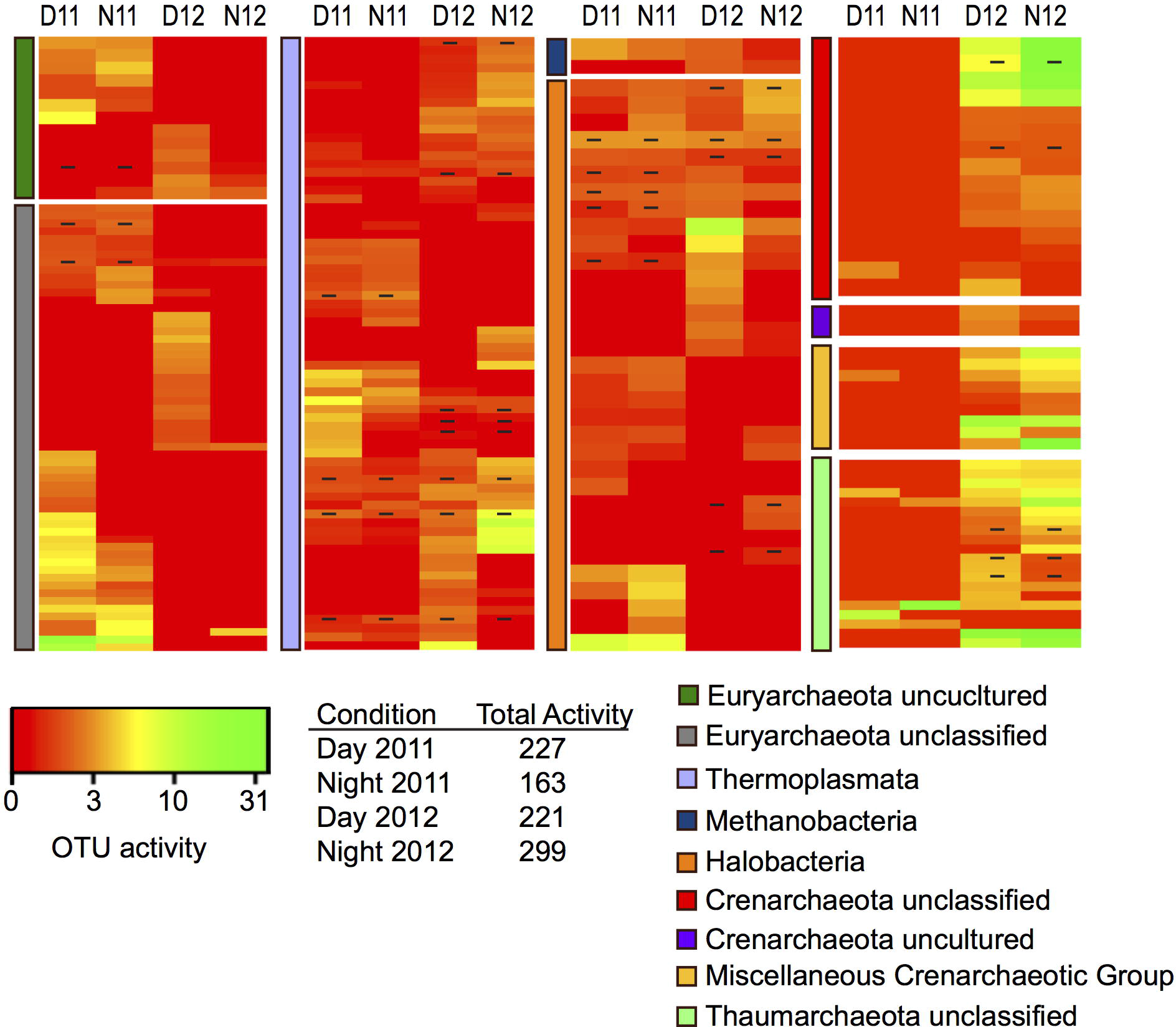
Heatmap analysis of OTU activity for each archaeal class for day and night samples across years (D11, Day 2011; N11, Night 2011, D12, Day 2012; N12, Night 2012). The heatmap color represents the extent of activity as measured by 16S rRNA:16S rRNA gene ratio for each OTU. Square colors shifted towards brighter green indicate higher activity of that OTU. Dash within squares indicates abundant OTUs. Table shows combined OTU activity (sum of all 16S rRNA:16S rRNA gene ratios) for each condition across years.

### Post-disturbance shifts in Bacterial OTU Activity

As opposed to archaeal OTUs, no significant association between activity and abundance was observed for bacterial OTUs during daytime across both salinity conditions, implying highly variable growth among bacterial taxa within the microbial mat ecosystem (Figure 2C; Kendall’s nonparametric τ_day2011_=0.05, p=0.15, n=434; τ_day2012_=0.06, p=0.05, n=697). However, a weak positive correlation was observed for nighttime samples across years (Figure 2D; Kendall’s nonparametric τ_night2011_=0.35, p<0.0001, n=347; τ_night2012_=0.26, p<0.0001, n=621). Analysis of bacterial OTUs during daytime showed that 12% of the abundant OTUs and 9% of the conditionally rare OTUs were active at high salinity (Figure 2C, red triangles). Following the reduction in salinity, a 35% decrease in active-abundant OTUs was observed coupled with a 60% increase in active-rare OTUs. Furthermore, nighttime samples showed that 24% of the abundant and 9% of the rare OTUs were active at high salinity (Figure 2D). Similar to other trends, post-disturbance conditions resulted in a 45% and 19% increase in active abundant and rare OTUs, respectively. These results are consistent with a previous study showing decreased bacterial diversity under high salinity conditions (Benlloch *et al.*, 2002) and suggest that the reduced salinity occurring after Hurricane Irene provided more favorable environmental conditions which resulted in overall increased bacterial activity.

Similar to Archaea, bacterial OTUs between and within phyla also demonstrated unique activity profiles across salinity and diel conditions (Figure 4). Examining pre-disturbance bacterial activity showed that the richness of the active bacterial community was low with most of the activity contained among unique OTUs within the Proteobacteria, Cyanobacteria, and Bacteroidetes phyla (Figure 4, D/N11). This activity is consistent with other studies that have found these phyla to contain halophilic and halotolerant taxa (Oren, 2008; Schneider *et al.*, 2013). Following the reduction in salinity, an increase in the richness of the active bacterial community was observed, resulting in increased activity across most phyla and an overall 69% increase in total bacterial activity as determined by the sum of day/night 16S rRNA:rRNA gene ratios for both salinity conditions (Figure 4, D/N12). Interestingly, while cyanobacterial OTUs increased in abundance following the salinity shift (Figure 1B), their overall activity decreased (Figure 4, D/N12). While the bacterial community across conditions showed differences in day/night activity, the trend was enhanced under the 2012 low salinity condition with OTUs within most phyla decreasing in activity at night, resulting in a 53% decrease in activity compared to daytime. The Proteobacteria phyla was further resolved to better understand how the salinity shift affected the activity of this metabolically-diverse group (Figure 5). Following salinity reduction, shifts in active OTUs within and across orders were observed, resulting in a 20% increase in overall proteobacterial activity. The Alphaproteobacteria was most abundant proteobacterial class across conditions (Figure 1B) and showed an 18% increase in overall activity under lower salinity (Figure 5, Alphaproteobacteria). Among the Alphaproteobacteria, increased activity was observed in OTUs classified within the nitrogen fixing Rhizobiales and the purple non-sulfur bacteria, Rhodobacterales and Rhodospirillales. Possibly due to having the highest OTU richness among proteobacterial classes (Figure 1B), the Deltaproteobacteria showed the greatest increase (40%) in activity with OTUs among all orders increasing following salinity reduction (Figure 5, Deltaproteobacteria). The Gammaproteobacteria had the greatest increase in abundance following salinity reduction due to the expansion of 3 OTUs contained within the Chromatiales and Sedimenticola orders (Figure 1B) which showed concomitant increases in activity (Figure 5, Gammaproteobacteria). Overall, the reduction in salinity resulted in increased bacterial activity with significant active/dormant shifts occurring among abundant and rare taxa that likely play a role in microbial mat biogeochemical cycling.

**Figure 4.**
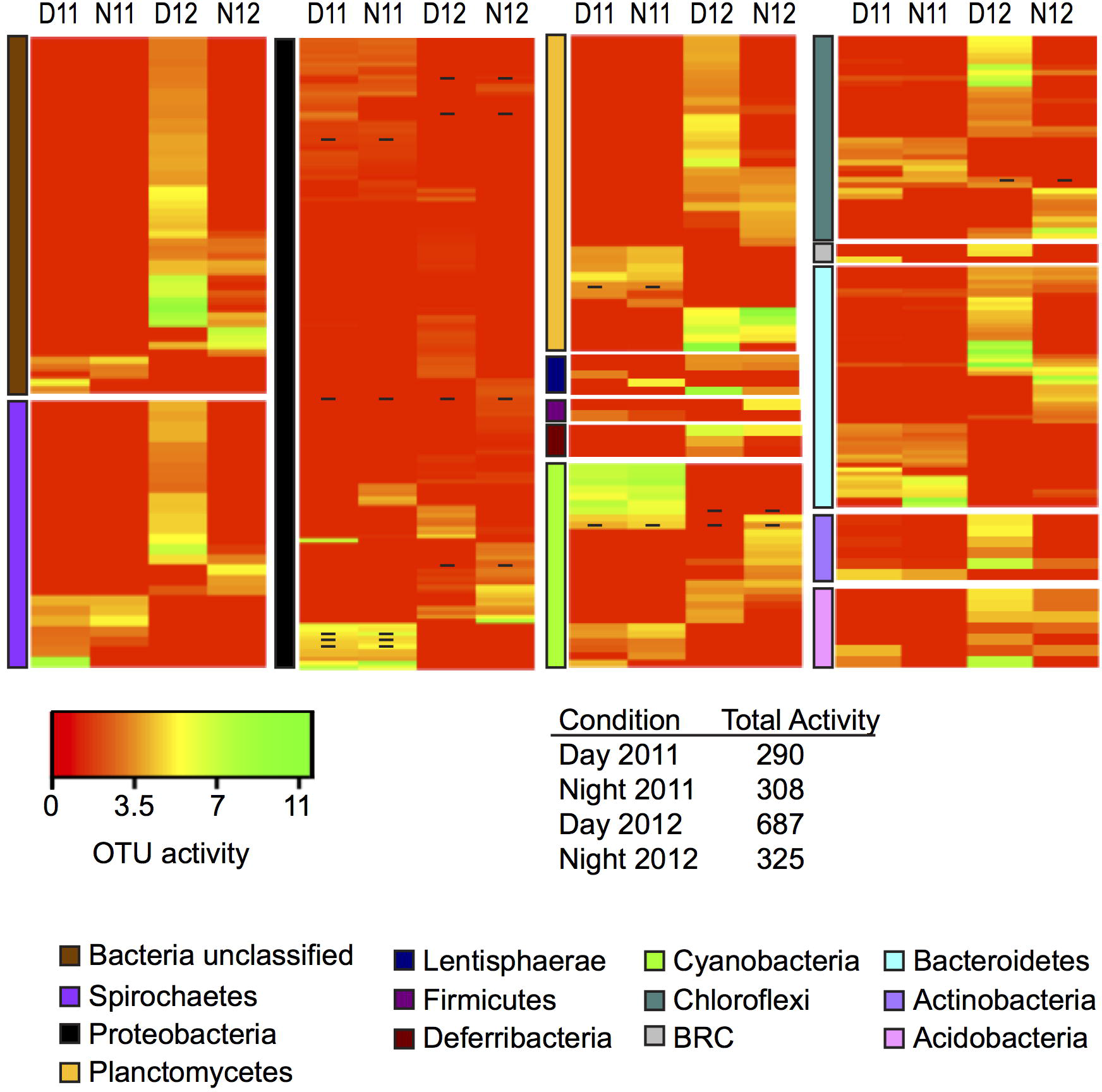
Heatmap analysis of OTU activity for each bacterial phylum for day and night samples across salinities (D11, Day 2011; N11, Night 2011; D12, Day 2012; N12 Night 2012). The heatmap color represents the extent of activity as measured by 16S rRNA:16S rRNA gene ratio within each OTU. Square colors shifted towards brighter green indicate higher activity of that OTU. Dash within squares indicates abundant OTUs. Table shows combined OTU activity (sum of all 16S rRNA:16S rRNA gene ratios) for each condition across years.

**Figure 5.**
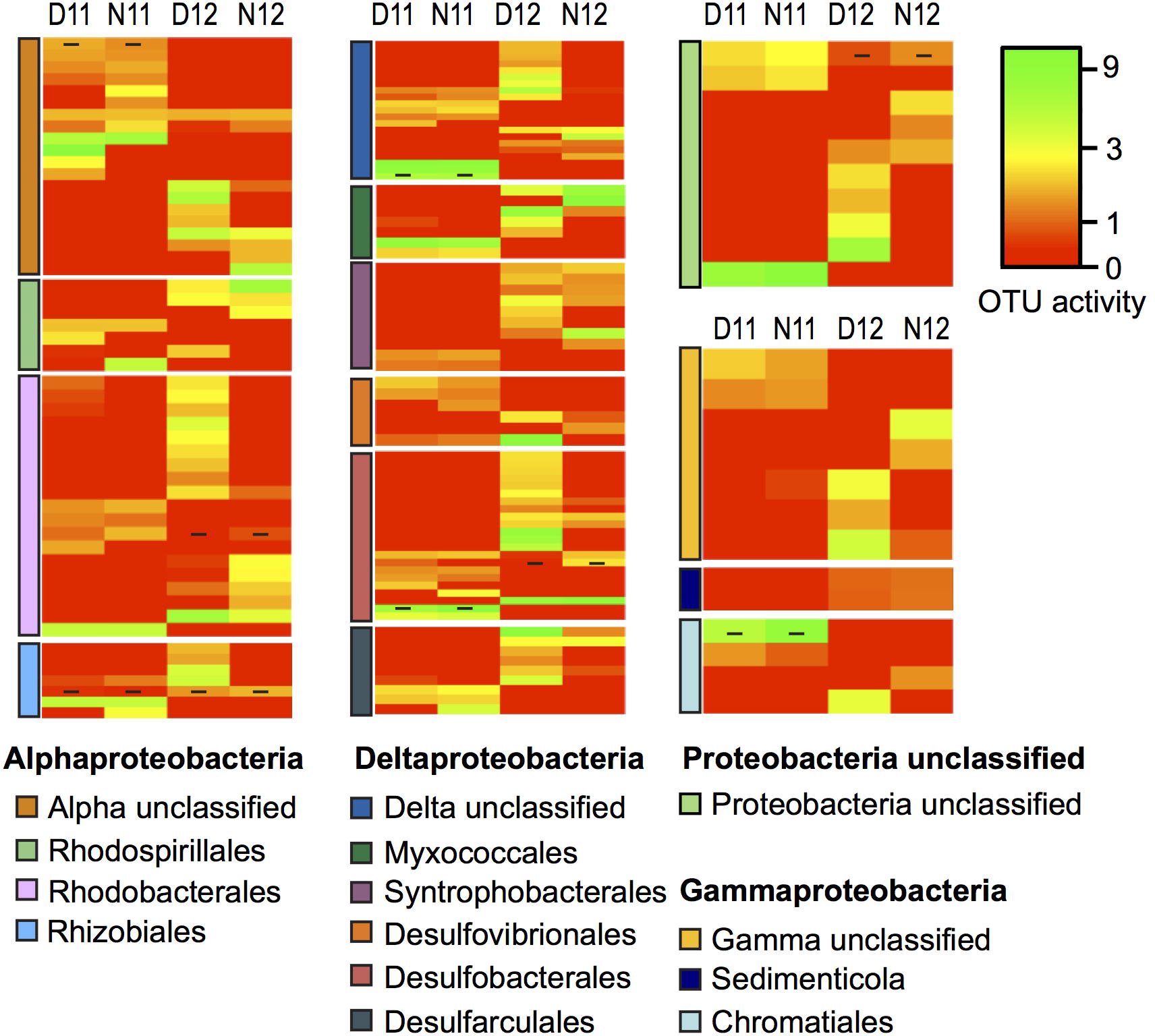
Heatmap analysis of OTU activity for each proteobacterial order for day and night samples across salinities (D11, Day 2011; N11, Night 2011; D12, Day 2012; N12 Night 2012). The heatmap color represents the extent of activity as measured by 16S rRNA:16S rRNA gene ratio within each OTU. Square colors shifted towards brighter green indicate higher activity of that OTU. Dash within squares indicates abundant OTUs.

### Biogeochemical cycling

Microbial mats are often considered natural bioreactors due to the close microbial metabolic coupling occurring within a millimeter scale and resulting in enhanced biogeochemical cycling (Visscher and Stolz, 2005). The shift in salinity experienced by the mat ecosystem investigated in this study following Hurricane Irene resulted in dynamic changes in the abundance and activity of archaeal and bacterial taxa. Many of these taxa likely play a role in biogeochemical processes; therefore, we next examined how the post-disturbance community shifts affected expression of nitrogen and sulfur cycling genes. Quantitative PCR was used to determine the copy number of nitrogen and sulfur cycling genes occurring within DNA and cDNA as a measure of genetic potential and expression, respectively (Figure 6). Other studies have used a similar approach to identify nitrogen and sulfur cycling potential and expression and have observed a correlation between gene expression and process rates (Church *et al.*, 2005; Philippot and Hallin, 2005; Bernhard *et al.*, 2010; Turk-Kubo et al., 2012; Akob *et al.*, 2012; Headd and Engel, 2013; Bowen *et al.*, 2014). However, it should be noted that potential post-transcriptional modifications and temporal decoupling (Chen *et al.*, 1998; Kim *et al.*, 1999; Nogales *et al.*, 2002; van de Leemput *et al.*, 2011) may confound the link between gene expression and process rates and is therefore used in this study as a comparative measure to assess how salinity influenced the biogeochemical potential within the microbial mat ecosystem.

**Figure 6.**
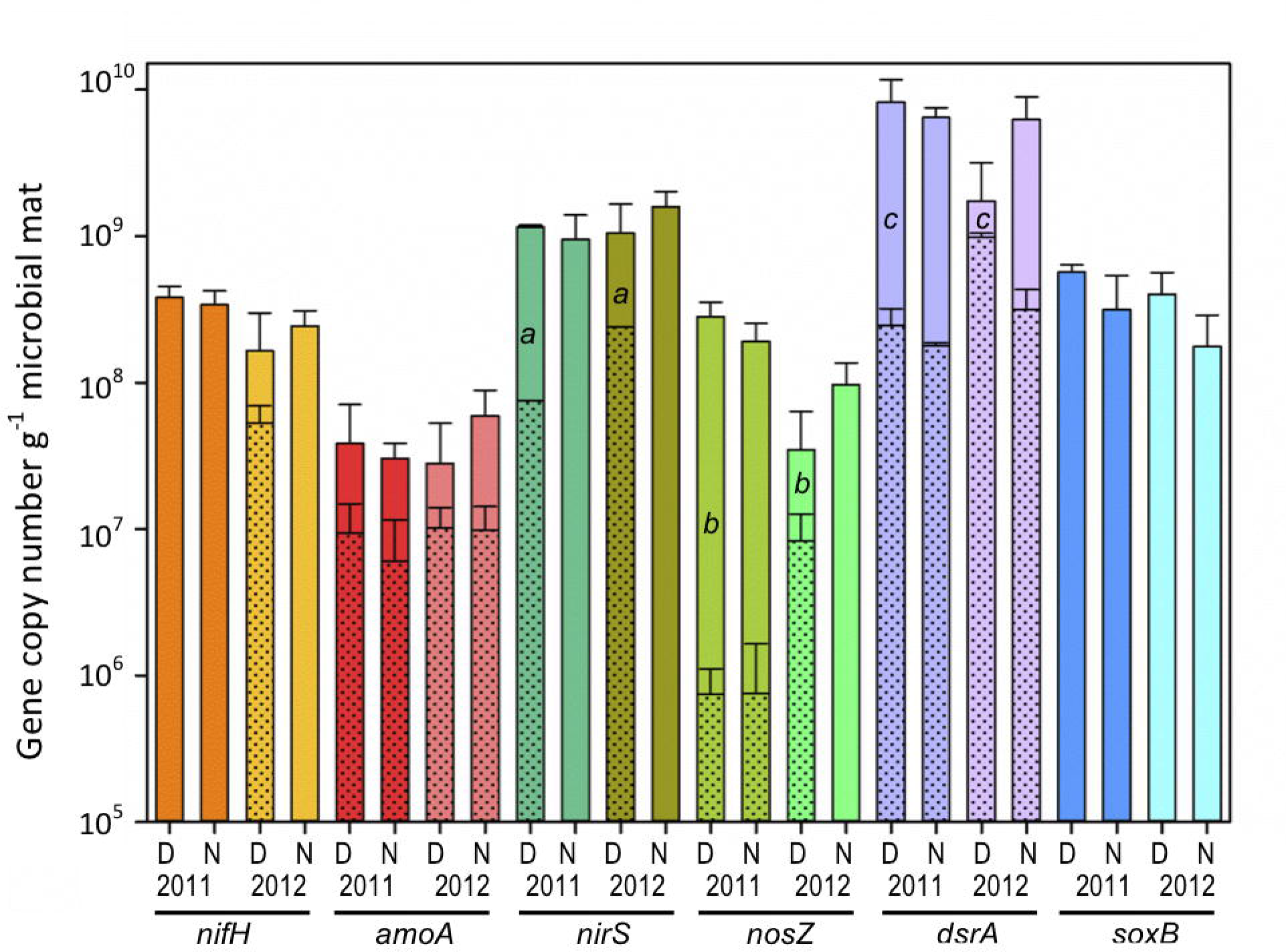
Microbial mat community potential for biogeochemical cycling as determined by qPCR amplification of genes (DNA) and transcripts (cDNA) involved in nitrogen and sulfur cycling for day (D) and night (N) samples during 2011 (11) and 2012 (12) (n=6). Nitrogen cycling genes/transcripts analyzed: nitrogen fixation (*nif*H), archaeal nitrification (*amoA*), and denitrification (*nosZ*, and *nirS*). Sulfur cycling genes/transcripts analyzed: sulfate reduction (*dsrA*) and sulfide oxidation (*soxB*). Shaded areas within solid bars represent transcript copy numbers as an estimate of gene expression (no shade indicates that transcripts were not detected). Asterisk indicates significant different cDNA/DNA ratios across years (independent t-test, p<0.05).

### Nitrogen fixation

The dinitrogenase reductase gene (*nifH*) was measured to examine community nitrogen fixation potential (Figure 6). Across both pre- and post-disturbance conditions, no significant difference (p<0.001) was observed in *nifH* DNA gene copy numbers per gram of microbial mat during day (2011: (3.8 ± 0.7)× 10^8^; 2012: (1.6 ± 1.4)x 10^8^) or night (2011: (3.4 ± 0.8) ×10^8^; 2012: (2.4 ± 0.7) x10^8^), suggesting overall community genetic potential for nitrogen fixation remains similar across salinities (Figure 6, *nifH* D/N11 vs. D/N12). However, using the cDNA/DNA ratio as a measure of potential *nifH* gene expression, an increase in daytime expression from below detection to 33% was observed following the reduction in salinity (Figure 6, *nifH* D11 vs D12 shaded bar). These data are consistent with a study observing suppression of N_2_ fixation in microbial mats at salinities over 90 g kg^-1^ (Pinckney *et al.*, 1995), and suggest that while nitrogen fixation potential was present, expression was repressed under the osmotic stress of extreme saline conditions. Diel studies of N_2_ fixation have observed complex light/dark regulation of *nifH* expression within different ecosystems with some microbial communities demonstrating increased expression during daytime and others enhanced under dark conditions (Pinckney *et al.*, 1995; Colón-López *et al.*, 1997; Church *et al.*, 2005; Severin and Stal, 2010; Woebken *et al.*, 2015). In this study, nighttime *nifH* expression was below detection under both salinities, suggesting that N_2_ fixation may be coupled to phototrophy through direct (*i.e.*, heterocystous Cyanobacteria) or indirect (*i.e.*, photosynthate utilization by noncyanobacterial diazotrophs) pathways. While the contribution of individual taxa to *nifH* expression was not established in this study, many OTUs within Classes/Orders known to contain diazotrophic members, such as Methanobacteria, Rhizobiales, and Desulfobacterales, demonstrated increased activity and abundance in daytime samples following the post-disturbance reduction in salinity.

### Nitrification

Bacterial and archaeal ammonia monooxygenase genes (*amoA*) were analyzed to investigate the potential contribution of nitrification among ammonia-oxidizing Archaea (AOA) relative to ammonia-oxidizing Bacteria (AOB). Attempts to amplify bacterial *amoA* genes from pre- and post-disturbance conditions were unsuccessful and typical AOB taxa were not identified in Proteobacteria 16S rRNA or rRNA genes, suggesting that bacterial ammonia oxidation is not present within this hypersaline ecosystem. However, archaeal *amoA* genes were identified in DNA at similar copy numbers with no significant difference (p>0.5) across years [2011: day (3.8 ± 3.2) x10^7^; night (3.0 ± 0.9) ×10^7^] and [2012: day (2.7 ± 2.5) ×10^7^); night (5.9 ± 6.7) ×10^7^], suggesting the role of the archaeal community in ammonia oxidation within this ecosystem (Figure 6, *amoA* D/N11 vs. D/N12). While AOB have been found in coastal microbial mats (Fan *et al.*, 2015), their absence within the Salt Pond ecosystem is not surprising as salinity has been shown to influence the distribution of ammonia oxidizers with AOA abundance correlated with increased salinity (Stehr *et al.*, 1995; Cébron *et al.*, 2003; Caffrey *et al.*, 2007; Beman *et al.*, 2008; Bernhard *et al.*, 2010; Lipsewers *et al.*, 2014). Analysis of *amoA* gene expression during day and night showed average cDNA/DNA ratios of 35% and 13% under high salinity conditions and increased to 63% and 20% following the 2012 post-disturbance salinity reduction. As discussed above (Figures 1 and 3), while taxa within the Thaumarchaeota phylum were active at day and night under both salinities, OTU abundance, richness, and activity shifted under the lower salinity condition and likely played a role in the increased *amoA* expression. In addition, given that AOA require molecular oxygen and show possible pH sensitivity (Beman *et al.*, 2011; Gubry-Rangin *et al.*, 2015), the observed differences in diel expression may result from partial physiological constraints as the photosynthetically-driven mat shifts from daytime oxic-alkaline conditions to nighttime anoxic-acidic conditions (Revsbech *et al.*, 1983). For instance, one Thaumarchaeota OTU further classified within the Nitrososphaeria class had increased activity during day versus night. Members of this class have been shown to have a preference to alkaline conditions (Gubry-Rangin *et al.*, 2015), suggesting shifts in pH may regulate the activity of this OTU. Thus, while archaeal *amoA* genes have been identified in a range of ecosystems (Francis *et al.*, 2005; Stahl and de la Torre, 2012; Yang *et al.*, 2013), the expression observed under the high salinity condition in this study expands the known range of salinity tolerance and suggests that the highly efficient AOA autotrophic pathway (Walker *et al.*, 2010; Konneke *et al.*, 2014) provides the energy necessary for survival and activity within hypersaline environments.

### Denitrification

The process of denitrification, reduction of nitrate or nitrite to nitrous oxide or dinitrogen, is the major mechanism by which fixed nitrogen returns to the atmosphere from soil and water. The denitrification potential of the community was estimated through analysis of the *nirK/nirS* (nitrite reductase) and *nosZ* (nitrous oxide reductase) genes. While both *nirK and nirS* have been identified in microbial mats across a range of salinities (Desnues *et al.*, 2007; Fan *et al.*, 2015), amplification of nitrite reductase genes from the Salt Pond microbial mat under both salinity conditions only identified *nirS* genes. This result supports other studies suggesting possible environmentally-driven differences in the ecological distribution of *nirK/nirS* genes (Oakley *et al.*, 2007; Smith and Ogram, 2008; Jones and Hallin, 2010). Within microbial mat DNA, similar (p>0.5) *nirS* and *nosZ* gene copy numbers were observed during day (2011: *nirS* (1.2 ± 0.03) x10^9^, *nosZ* (2.8 ± 0.7) x10^8^; and 2012: *nirS* (1.1 ± 0.6) x10^9^, *nosZ* (3.5 ± 2.8) x10^7^) and night (2011: *nirS* (9.5 ± 2.8) x10^8^, *nosZ* (1.9 ± 0.6) x10^8^; and 2012: *nirS* (1.6 ± 0.2) ×10^9^, *nosZ* (9.7 ± 3.4) =10^7^) under both salinity conditions, suggesting the overall denitrification potential of the community remained similar in both pre- and postdisturbance conditions (Figure 6). However, comparison of *nirS* to *nosZ* across both years indicated that the abundance of *nirS* was greater than *nosZ*. This supports another study that observed decreased gene abundance along the denitrification pathway (Bru *et al.*, 2011), which is likely linked to the sequential reduction in energy gained by each step (Koike and Hattori, 1975). While *nirS* and *nosZ* gene abundance remained similar across years, significant increases (p<0.001) in daytime cDNA/DNA ratios for both genes were observed during 2012 (Figure 6, *nirS, nosZ:* D11 and D12), suggesting increased gene expression and potential for denitrification with reduced salinity. While rates of denitrification are regulated by many variables (*e.g.*, oxygen concentration, pH, temperature, salinity), the greatest influence is often considered to be carbon and nitrogen availability (Hallin *et al.*, 2009; Jones and Hallin, 2010; Babbin and Ward, 2013). Extreme salinities have been shown to reduce microbial mat photosynthetic carbon fixation and nitrogen fixation; therefore, the postdisturbance reduction in salinity likely increased carbon and nitrogen availability and enhanced denitrification. When examining potential differences in diel expression, *nirS* cDNA was below detection at night in both years while *nosZ* showed minimal (2.2%) nighttime expression during 2011 and below detection in 2012. This suggests that denitrification might be coupled to anoxygenic photosynthetic bacteria as has been observed in lake sediments (Shen and Hirayama, 1991) or that the routine diel changes in pH, oxygen, and carbon availability provide an environmental constraint that limits denitrification to daytime. A range of taxonomically diverse heterotrophic and autotrophic microbes are capable of denitrification, making phylogenetic inference difficult (Zumft, 1997; Bowen *et al.*, 2014). However, it is known that halophilic archaea within the phyla Euryarchaeota and some taxa within the alphaproteobacterial orders, Rhodospirllales and Rhodobacterales, play a role in denitrification (Tomlinson *et al.*, 1986; Yoshimatsu *et al.*, 2000; Hattori *et al.*, 2016). While taxonomic links to microbial mat denitrification are speculative, OTUs classified within the Halobacteriaceae, Rhodospirllales, and Rhodobacterales were active across years but increased activity following salinity reduction, and may play a role in denitrification within this ecosystem. Anaerobic ammonia oxidation (anammox) was also investigated and while taxa affiliated with the Plantomycetes phylum were identified, attempts to amplify the hydrazine oxidoreductase (hzo) genes were unsuccessful.

### Sulfate reduction

Microbial mats typically have high rates of primary productivity resulting in the production and excretion of photosynthates into surrounding sediments (Bateson and Ward, 1988). Within marine hypersaline systems, these organic compounds are mineralized, in large part, by microbes using anaerobic respiration coupled to sulfate reduction (Baumgartner *et al.*, 2006; Brocke *et al.*, 2015). In this study, the dissimilatory sulfite reductase (*dsrA*) gene was analyzed to estimate pre- and post-disturbance sulfate reduction potential and expression by the Salt Pond microbial mat community (Figure 6). There were no significant differences (p>0.5) in *dsrA* gene abundance in day/night DNA during both years (day11: (8.2 ± 2.0) x10^9^, night11: (6.4 ± 1.1) x10^9^; day12: (1.7 ± 1.4) x10^9^), night12: (6.2 ± 2.7)x10^9^), suggesting that the community retains similar sulfate reduction potential under both high and low salinity conditions. While gene abundance was equal across conditions, cDNA/DNA ratios indicated low-level *dsrA* expression (≤7%) under all conditions except in the 2012 lower salinity day samples. Under the 2012 day condition, the average cDNA/DNA ratio increased to 81% representing a significant increase (407.7%, p<0.001) in expression and suggesting that sulfate reduction may have been suppressed by high salinity conditions until the postdisturbance reduction in salinity lowered the environmental constraint resulting in increased *dsrA* gene expression and possibly sulfate reduction within this ecosystem. Others have also observed decreased sulfate reduction at salt saturation (Oren, 1999; Kulp *et al.*, 2007; Gu *et al.*, 2012). While the exact mechanism of suppression is unclear, it has been hypothesized that sulfate reduction may not supply enough energy for osmoadaptation (Oren, 1999, 2011) or that increased borate concentration occurring at salt saturation may act as a specific inhibitor of this metabolism (Kulp *et al.*, 2007). When examining day/night differences in 2012, cDNA/DNA ratios indicate that *dsrA* expression was greatest during the day and suppressed at night. While sulfate reduction is considered to be a strict anaerobic process, oxygen tolerance has been observed (Cypionka, 1995) and provides a selective advantage by increasing accessibility to carbon derived from photosynthetic processes. This finding is in agreement with other studies that observed high rates of sulfate reduction in the upper layers of microbial mats coupled to carbon fixation by the phototrophic community (Teske *et al.*, 1998). The ability to use sulfur species as an electron acceptor has been identified in a taxonomically diverse group of Archaea and Bacteria. For instance, within the Archaea, all known sulfate reducing taxa are considered hyperthermophiles and have been classified within the Crenarchaeota, Thermoplasmata, and Methanobacteria (Liu *et al.*, 2012). Within Bacteria, sulfate reducing taxa are mostly located within the Deltaproteobacteria and Epsilonproteobacteria and have been identified in a wide range of ecosystems, including microbial mats (Muyzer and Stams, 2008). Under the 2011 high salinity condition, while all potential sulfate reducing groups were active, taxa within the Thermoplasmata had the highest OTU richness and abundance, suggesting these taxa may play a greater role in maintaining the sulfur cycle under extreme salinity (Figures 1, 3, 5). Following the postdisturbance reduction in salinity, OTU richness, abundance, and activity increased for all groups concomitant with significant increases in *dsrA* expression. However, OTUs within the Deltaproteobacteria showed the greatest increase in richness and activity during daytime, suggesting a possible shift in the predominant sulfate reducing group. Overall, it is likely that within this microbial mat ecosystem, salinity influences the major sulfate reducing taxa with archaeal Thermoplasmata taxa playing a greater role under higher salinities and bacterial Deltaproteobacteria playing a greater role in lower salinity conditions.

### Sulfide oxidation

At zones within microbial mat ecosystems at which oxygen and sulfide overlap, sulfate reduction is closely coupled to the oxidation of reduced sulfur compounds by chemolithotrophic and anoxygenic phototrophic microorganisms utilizing the Sox enzymatic system (Headd and Engel, 2013). Among the Sox system, the *soxB* gene, encoding the periplasmic thiosulfate-oxidizing Sox enzyme, is often used as a marker of microbial community sulfide oxidation potential (Headd and Engel, 2013). In this study, we measured the abundance of the *soxB* gene to determine the sulfide oxidizing potential and activity by the Salt Pond community during day and night under high and low salinity. There were no significant differences in *soxB* gene copy numbers in DNA extracted from day (2011: (5.7 ± 0.7) x 10^8^; 2012: (4.0 ± 1.6) x 10^8^) and night (2011: 3.1 ± 2.2) x 10^8^; 2012: (1.8 ± 1.1) x 10^8^) samples across years, suggesting the potential for sulfide oxidation exists under both salinity conditions (Figure 6). However, no *soxB* genes were detected in cDNA isolated from the same samples (other biogeochemical genes were amplified from these samples), suggesting that while sulfide oxidation potential exists, the gene is suppressed under all analyzed conditions. The mechanism of this suppression is unclear and is the focus of future studies. However, it is possible that, while sulfide oxidation is thermodynamically favorable for osmoadaptation, increased levels of salinity or other environmental factors may have suppressed the activity of the sulfide oxidizing community. Evidence for a possible salinity-driven suppression of sulfide oxidation was observed in a study examining the sulfur cycle along a salinity gradient from freshwater into the Dead Sea (Häusler *et al.*, 2014). This study observed that increasing salinity selected for different sulfide oxidizing communities with the highest salinity restricting growth to only halotolerant or halophilic taxa. Similarly, while OTUs affiliated with possible sulfide oxidizing groups such as the acidophilic archaea Thermoplasmatales (Euryarchaeota) and Chromatiales were detected in Salt Pond samples, their activity was low across both salinity conditions. Therefore, the sulfide oxidizing community within this ecosystem may be composed of taxa adapted to lower salinity conditions resulting in the suppression of *soxB* expression.

### Conclusion

The microbial mat examined in this study is located within a coastal lagoon ecosystem that experiences routine but wide ranging seasonal variation in environmental conditions. The extensive variability of this system may select for a highly diverse microbial community that provides ecosystem stability following weather-driven disturbances. The ecosystem-level pulse disturbance generated by Hurricane Irene significantly reduced local salinity by 72%, subsequently resulting in Archaea to Bacterial domain shifts. Detailed analysis of this broader trend showed significant shifts among taxa within rare and abundant archaeal and bacterial populations, with less than 1% of taxa remaining abundant. In both archaeal and bacterial communities, the majority of taxa were slow-growing/dormant across salinities; however, post-disturbance samples showed that the total number of active taxa and overall activity increased with significant active/dormant shifts occurring among abundant and rare taxa across and within phyla. While changes in the abundance and activity of specific taxa can not be linked to individual biogeochemical processes, overall post-disturbance shifts in community structure enhanced expression of genes involved in N and S cycling. Together, these data show the functional and compositional sensitivity of a microbial mat ecosystem to environmental change but also suggest that rare taxa provide a reservoir of genetic diversity that may enhance ecosystem stability following seasonal and extreme environmental disturbances. While this study begins to address the complex links between microbial diversity and ecosystem processes, more detailed studies are currently underway and aimed at further defining the ecological principles examined in this study.

## Conflict of interest

The authors declare no conflict of interest.

## Authors and contributors

Microbial mat samples were collected and processed by E.C.P and R.S.N. Laboratory sample processing pipeline was developed by E.J.B and E.C.P. Statistical and computational analysis was performed by E.C.P and R.S.N. Manuscript was reviewed and edited by all authors.

## Funding

Funding for this study was provided by the National Science Foundation (RSN, grant numbers: EF-0723707 & DEB-1149447) and further computational support was provided by the XSEDE program (RSN, grant number TG-DEB140011).

## Acknowledgements

We would like to thank the staff at the Gerace Research Centre on San Salvador Island, The Bahamas for logistical support for this research. We also acknowledge the Research Cyberinfrastructure staff at the University of South Carolina for computational support.

